# How does Gloger’s rule predict near- and mid-infrared radiation in birds?

**DOI:** 10.1101/2024.12.12.628255

**Authors:** Thomas Lee, Meghan Barrett, Laurent Pilon, Allison J. Shultz, Terrence P McGlynn

## Abstract

Animal coloration has diverse functions such as camouflage, communication, thermoregulation, protection from UV damage and more, and can be shaped by environmental selective pressures. Some climactic selective pressures are strong enough to produce consistent patterns in many species across large-scale geographic gradients. One pattern in endothermic animals is Gloger’s rule, which predicts that populations in hot, humid areas will be darker than those in cool, dry areas. This pattern has been demonstrated in several species across latitudinal gradients and is expected to relate to the selective effects of both local thermoregulatory pressures and humidity. However, shortwave radiation from sunlight extends beyond the visible spectrum [400-700 nm] into the near-infrared; thus, thermal pressures often result in changes in surface reflectance characteristics beyond the visible [e.g., 700-2500 nm]. Further, heat exchange with the environment extends into the mid-infrared, including MIR heat loss through the atmospheric transmission window [7.5 - 14 um]. Here, we examine both UV-NIR absorption and MIR emittance in five species of birds that have been shown to follow, or not follow, Gloger’s rule. We show that NIR absorption varies by species and population in ways that correspond to their habitat and thermoregulatory strategies. MIR emittance, by contrast, was very stable across both species and populations but differed across populations of Northern Bobwhites. We conclude by highlighting the importance of extending coloration research into the NIR and MIR. Further consideration of infrared radiation is necessary for a complete view of animals’ phenotypic diversity and possible responses to thermal challenge.

## Introduction

External coloration is one mechanism by which animals adapt to various selective pressures in their environment. In the visible [400 - 700 nm] spectrum, this complex phenotypic trait has diverse functions, such as camouflage, intraspecific communication, parasite deterrence, thermoregulation, and more (Cuthill et al. 2017; Stuart-Fox et al. 2017). The ultraviolet through near infrared [300 - 2500 nm] and mid-infrared [MIR; 2.5 - 20 μm] wavelengths of light also have thermoregulatory functions corresponding to absorptance of the solar irradiance and the emittance of the organism at its average core temperature (Stuart-Fox et al. 2017; Krishna et al. 2020). Where coloration is expected to have a thermoregulatory function, climate variation across large-scale geographic gradients is often correlated with animal coloration (Chown et al. 2004).

Gloger’s rule is a commonly-observed pattern describing the relationship between climate and color in birds and mammals . The simple version of this rule posits that endothermic animals will be darker in humid and warm areas due to the increased deposition of melanin in their surface structures (Delhey 2019). This macroecological pattern is often observed across latitudinal gradients, with darker populations of the same species found in wetter and warmer environments closer to the equator (e.g., Delhey et al. 2019). Although temperature and humidity have traditionally both been expected to be important for Gloger’s rule, humidity has a stronger effect on melanin deposition and thus visible coloration after controlling for habitat type (Delhey 2019; Marcondes et al. 2021). Some have even proposed to redefine Gloger’s rule only in terms of humidity (Delhey et al. 2020).

If Gloger’s rule is driven by humidity, then relationships observed in the visible spectrum may differ in the near-infrared, which is more strongly tied to thermoregulation. About 55% of incoming solar energy falls within the near infrared (NIR,wavelengths from 701 - 2500 nm; (Stuart-Fox et al. 2017)]. Thus, adaptations that change in the balance of an absorbance and reflectance in the NIR can play a strong role in thermoregulation. NIR reflectance cannot be seen by conspecifics or predators, and has not been shown to assist in protection against parasites or microbes (Stuart-Fox et al. 2017). Variation in NIR reflectance has been found to correlate strongly with climate variation in insects and birds (Medina et al. 2018; Munro et al. 2019; Kang et al. 2021; Mason et al. 2023; though see Porter et al. 2023; Wang et al. 2023).

In addition to the reflection of solar irradiance in the UV-NIR, animals can also exchange longwave radiation (MIR) with their environment as another mechanism of thermoregulation. Substantial heat emittance in the MIR (typically, 7.5 - 14 um; Howell et al. 2015) can allow animals to cool down by using space as a heat sink, due to the highly transmissive atmosphere across this spectral range (Taylor and Yates 1957). Alternatively, low emittance in the MIR wavelengths would allow animals to retain heat. Insects found in hot environments have increased emittance as a method of remaining cool, irrespective of other climate variables like precipitation and humidity (Krishna et al. 2020, 2021). Variation in MIR emittance can even be specific to a body region: for instance, the Saharan silver ants, known for foraging in an extremely hot microclimate, have very high emittance on their dorsal surface to lose heat via the atmospheric transmission window and low emittance on the ventral surface to avoid heat exchange with the superheated ground (Shi et al. 2015). However, endotherms like birds have different challenges in thermoregulating via longwave radiative heat exchange, and to our knowledge, macroecological relationships of MIR emittance in birds remain unexplored.

In this study, we test whether patterns consistent with Gloger’s rule appear in the NIR and MIR for endotherms. We selected three, geographically diverse populations from five species of birds with wide latitudinal distributions - four known to follow Gloger’s rule (Great Horned Owl *Bubo virginianus* , Northern Bobwhite *Colinus virginianus* , Song Sparrow *Melospiza melodia* , Steller’s Jay *Cyanocitta stelleri* ), and one not known to follow Gloger’s rule (Common Raven *Corvus corax* ). We also selected species that are known to be primarily open-habitat species (Northern Bobwhite, Common Raven), closed-habitat species (Song Sparrow, Steller’s Jay), or nocturnal species (Great Horned Owl), to facilitate comparisons among species with different solar exposure. We used spectrometry to assess absorption and emittance on the dorsal surface birds (the mantle) of each population from each species. First, we assessed whether populations followed Gloger’s rule in each species by looking at bird-visible (UV-inclusive, 300 - 700 nm) and human-visible (400 - 700 nm) absorption coefficients. Then, we assessed whether UV-NIR solar absorption and MIR emittance differ across populations in a manner consistent with Gloger’s rule for each species.

## Materials and Methods

### Sampling

For each species, we selected three populations with differences in environmental conditions, largely corresponding with latitudinal variation. From each population, we selected three adult specimens to account for individual variation (Supplemental Table 1). All individuals are from the Ornithology Department of the Natural History Museum of Los Angeles County.

### Climate Data Acquisition

Climate data was obtained from the NOAA National Centers for Environmental Information’s (NCEI) Annual Climate Maps tool, using the most recent data from the closest NOAA stations that contained temperature and precipitation data. Data was accessed from the interactive map at https://www.ncei.noaa.gov/maps/annual/ on November 8, 2024.

### Spectrometry

#### Normal-normal Reflectance Probe

We measured normal-normal reflectance in the spectral range of 300 - 700 nm using a UV-Vis spectrophotometer (Flame-S-UV-Vis, Ocean Optics), using a pulsed Xenon lamp (PX-2, Ocean Optics), and 400 μm reflectance probe (WS-1-SL, Ocean Optics) fitted with a modified rubber stopper to exclude all incident light. Reflectance spectra were collected relative to a spectralon diffuse white standard (WS-1-SL, Ocean Optics) with a 500 ms integration time, boxcar width of 5 and averaging 10 scans. To account for measurement uncertainty, we averaged three measurements for each patch measured from the dorsal surface of each specimen.

#### UV-Vis Spectrometer

The spectral normal-hemispherical reflectance R_nh,λ_ of the dorsal surface in the spectral range of 300 - 1100 nm was measured using a UV–Vis spectrometer (Evolution TM 201 UV-Visible Spectrometer, Thermo Scientific) fitted with an integrating sphere accessory (DRA-EV-600, Thermo Scientific). Integration time was set to 0.2 seconds with spectral increments of 1 nm. The spectral normal-hemispherical reflectance R_nh,λ_ was estimated according to:

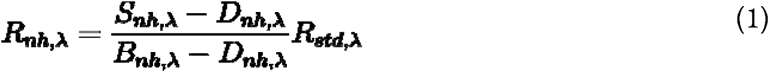

The spectral normal-hemispherical reflectance signal S_nh,λ_ measured by the UV-Vis spectrometer was corrected by subtracting the dark signal D_nh,λ_ measured by blocking any light from entering the detector. The baseline spectral normal-hemispherical reflectance measurement B_nh,λ_ was determined using a calibrated specular reflection standard mirror (NIST certified STAN-SSH, Ocean Optics) with known standard normal-hemispherical reflectance R_std,λ_.

#### FTIR

A nitrogen-purged Fourier transform infrared (FTIR) spectrometer (Nicolet TM iS50, Thermo Scientific Fischer, USA) equipped with an integrating sphere (Upward IntegratIRTM, PIKE Technologies, USA) was used to measure the spectral normal-hemispherical reflectance R_nh,λ_ of the dorsal surface in the NIR and MIR. A liquid-nitrogen cooled Mercury-Cadmium-Telluride (MCT) detector and a KBr beamsplitter were used to measure R_nh,λ_ in the range between 2 and 20 μm, and an InGaAs detector and a Calcium fluoride (CaF_2_) were used to measure R_nh,λ_ in the range between 1 and 2.5 μm. Similar to the UV-Vis spectrometer, the spectral normal-hemispherical reflectance R_nh,λ_ was estimated according to:

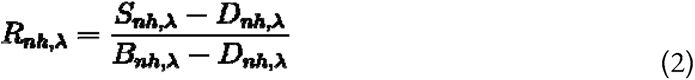

The spectral normal-hemispherical reflectance signal S_nh,λ_ measured by the FTIR was corrected by subtracting the dark signal D_nh,λ_ measured by blocking any light from entering the detector. The baseline spectral normal-hemispherical reflectance measurement B_nh,λ_ was determined using a gold reflectance standard (Thermofisher).

#### Parameters

To ensure reliable data collection between the two instruments, each bird was measured in the same local area in the center between the wings of the bird, and then the measurements (per instrument within the same spectral range) were averaged to minimize small variations in sampling area.

#### Process notes

For each bird species, the reflectance, dark, and baseline signals for a given spectrometer and detector ensemble was collected on the same day. For the owl and raven species, measurements on the FTIR were supported by an optical stand so that the region between the feathers could be measured consistently.

#### Post-collection data cleaning/manipulation information

Some of the spectral ranges measured by the spectrophotometers overlapped and to calculate total emittance or total absorptance, the data was condensed to have one reflectance value per wavelength. For the region between 1 and 1.1 μm, data sets collected from the UV-Vis and the InGaAs detectors were approximately linear in this region so a linear interpolation was used to merge both data sets. For the region between 2 and 2.5 μm, the MCT and InGaAs detectors captured similar reflectance curves indicative of chemical absorption peaks by N-H bonds in keratin (Miyamae et al. 2007). To preserve the shape of these curves, the MCT data was used for data processing so that the 2 to 20 μm could be kept continuously without any manipulation of data (Supplemental Figure 1 for example in sparrows).

The bird feathers are assumed to be dielectric materials that have optically rough surfaces so the emittance of the bird feathers are independent of direction (i.e., a diffuse emitter). Using Kirchoff’s law, the spectral normal-hemispherical emittance ε_nh,λ_ and spectral normal-hemispherical absorptance α_nh,λ_ is expressed as (Howell et al, 2015):

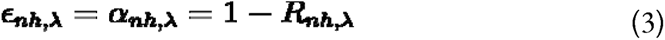

Then, the total normal-hemispherical emittance ε_nh_ is defined as (Howell et al, 2015):

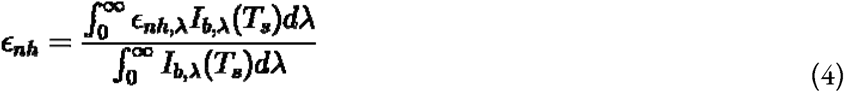

where I_b,λ_(T_s_) is the spectral Planck’s blackbody intensity at surface temperature, T_s_, taken as 313 K (Simpson, 1922), and integrals in the numerator and denominator were truncated with bounds of 0.3 μm and 20 μm for the available spectral data obtained by our instruments. This truncation considers 78% of full blackbody emission energy.

The solar absorptance αs was calculated according to:

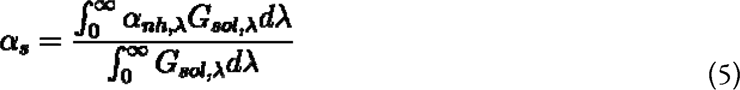

where G_sol,λ_ is the ASTM G-173 Spectra (Howell et al., 2015). Here also, the integrals were truncated with bounds of 0.3 μm and 20 μm for the available spectral data obtained by our instruments. This truncation considers 97% of the total blackbody radiation I_b_(T_s_) and solar radiation through AM1.5G. Additionally, we used the reflectance data to calculate the average brilliance of the birds (Trigo et al. 2015) using the formula shown below:

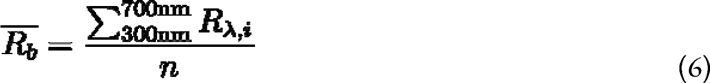

Where R_.o,i_ is the reflectance data at a wavelength between 300 and 700 nm and n is the number of integer wavelengths between 300 and 700 nm.

### Statistical Analyses

All data are available on Dryad: https://doi.org/10.5061/dryad.fxpnvx13b

We used GraphPad Prism v. 9.3.1 (GraphPad Software, 2023) to analyze all data. We used Shapiro-Wilk normality tests to assess normal distribution of the data. We used a one-way ANOVA followed by Tukey’s Multiple Comparisons Test for normally distributed data or a Kruskal-Wallis test with Dunn’s Multiple Comparisons Test for data that were not normally distributed. These methods were used to assess species-level differences in emittance and population-level differences in absorptance and emittance within each species. For ravens, we also tested differences between the two named subspecies by combining the data for two populations (for all other species all populations come from different named subspecies). Here, we used a Welch’s t-test to test for differences among subspecies in absorptance, to account for unequal variance, and an unpaired t-test to test for differences between subspecies in emittance. Normal-normal reflectance data was used for cross-species comparisons, and for all intraspecific comparisons with owls and ravens (as normal-hemispherical data could not be gathered for these species due to size-related limitations). Normal-hemispherical reflectance data was used for intraspecific comparisons in bobwhites, jays, and sparrows. Generalized linear models were used to assess the relationship between an individual’s UV-NIR reflectance and

For all tests, alpha was set to 0.05.

## Results

### Species-level absorptance and emittance

Normal-normal reflectance data demonstrated significant variation in mean solar absorptance (α) across species (Table 2, Figures 1 and 2; Kruskal-Wallis, K-W = 34.88, p < 0.0001). Owls had a significantly lower mean absorptance than ravens and sparrows (Dunn’s MCT: owl-raven: Z = 5.38, p <0.0001; owl-sparrow: Z = 3.27, p = 0.0109). Bobwhites also had lower mean absorption than ravens (Z = 4.38, p = 0.0001). There was no difference in the normal-hemispherical reflectance/absorption of bobwhites, sparrows, or jays (Kruskal-Wallis; K-W: 4.98, p = 0.0829).

**Figure 1.**
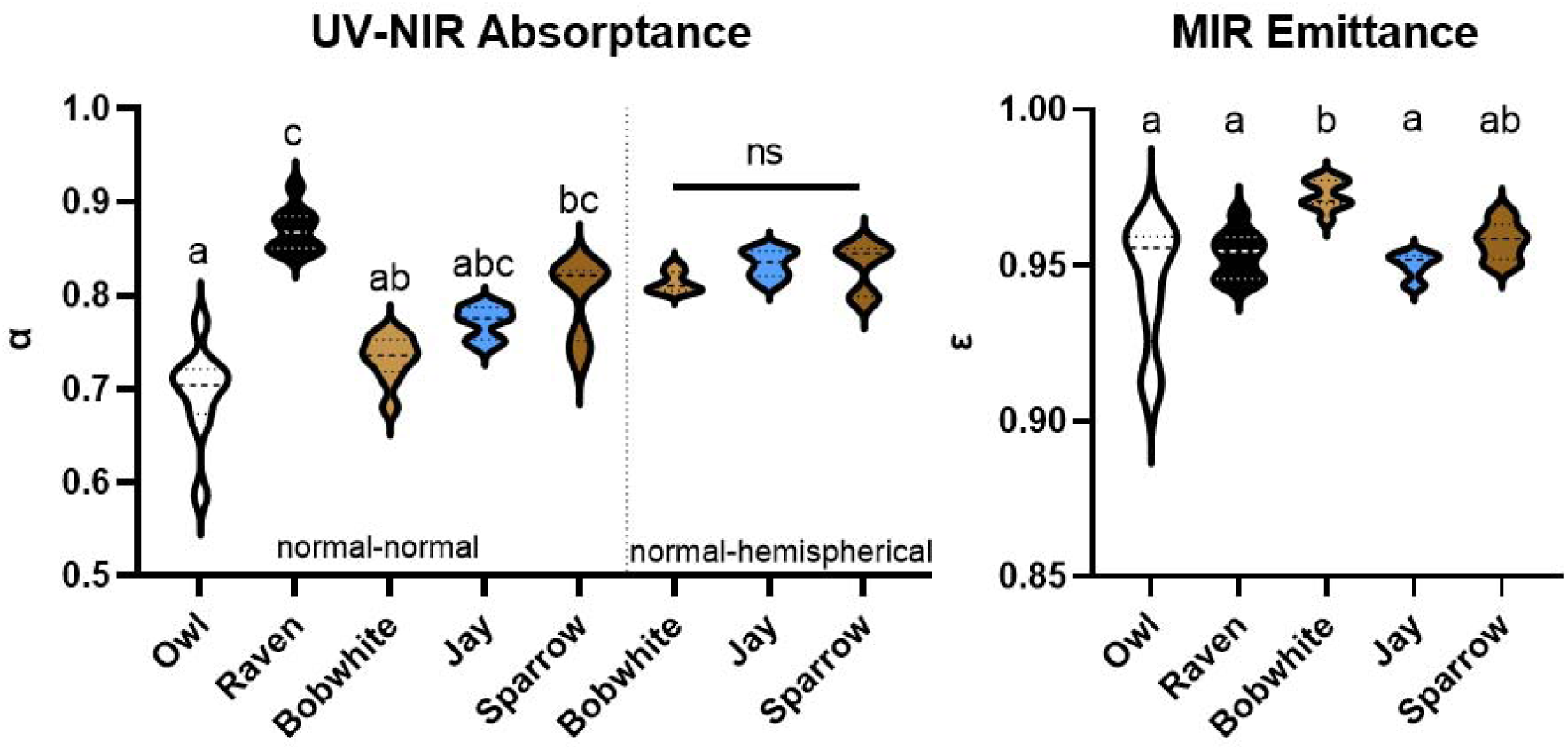
Variation in (left) solar absorptance and (right) emittance coefficients for each species. Normal-normal data demonstrated that there was significant variation in mean solar absorptance (α) across species (Table 2; Kruskal-Wallis, K-W = 34.88, p < 0.0001). Owls had a significantly lower mean absorptance than ravens and sparrows (Dunn’s MCT: owl-raven: Z = 5.38, p <0.0001; owl-sparrow: Z = 3.27, p = 0.0109); bobwhites also had lower mean absorptance than ravens (Z = 4.38, p = 0.0001). There was no difference in the normal-hemispherical reflectance/absorptance of bobwhites, sparrows, or jays (Kruskal-Wallis; K-W: 4.98, p = 0.0829). Bobwhites had significantly higher emittance than all species except sparrows (Dunn’s MCT; bobwhite-owl: Z = 3.89, p = 0.001; bobwhite-raven: Z = 3.46, p = 0.0053; bobwhite-jay: Z = 4.44, p < 0.0001); all other species were not significantly different (Z < 2.48, p > 0.05). Letters indicate statistically significant differences among species. ns = not significant.

**Figure 2.**
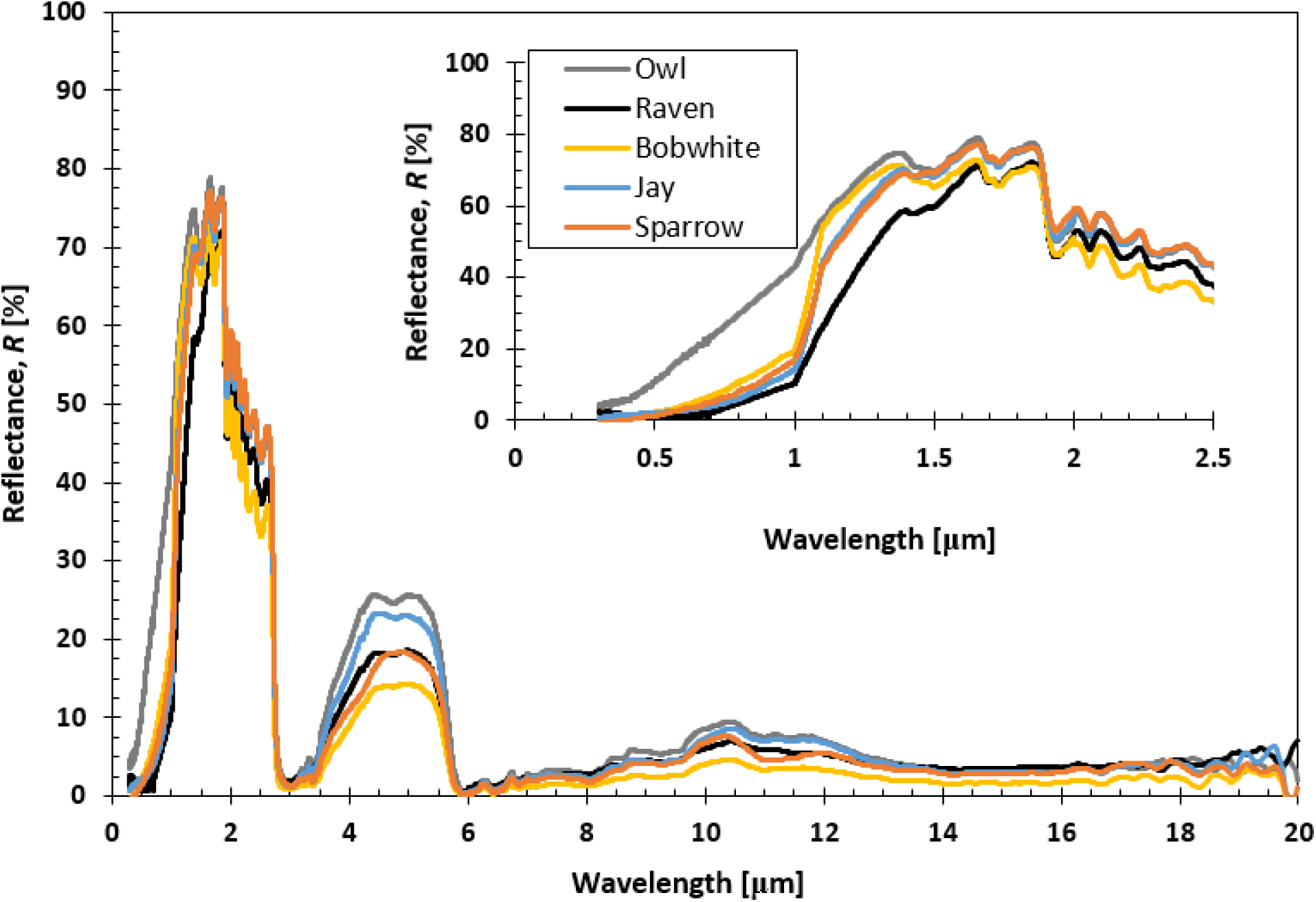
Average UV-NIR (normal-normal) and MIR reflectance spectra for each species of bird studied. Reflectance over the full spectra (0.3 - 20 um) is shown for each species with a solid line; the 95% confidence interval for each species is a dotted line. The inset shows the solar-relevant spectra (300 nm - 2500 nm, UV-NIR, normal-normal) in greater detail.

**Table 1:** Terms and Definitions.

| Table 1: Terms and Definitions |  |
| --- | --- |
| UV | Ultraviolet wavelengths of the electromagnetic spectrum, 300 nm<br>to 400 nm |
| VIS | Visible wavelengths of the electromagnetic spectrum, 400 nm to 700 nm |
| NIR | Near Infrared wavelengths of the electromagnetic spectrum, 0.7 $\mu\text{m}$ to 2.5 $\mu\text{m}$ |
| MIR | Mid Infrared wavelengths of the electromagnetic spectrum, 2.5 $\mu\text{m}$ to 20 $\mu\text{m}$ |
| Longwave Radiation | Radiative exchange associated with longer wavelengths that are emitted and absorbed by a surface; wavelength range includes the MIR |
| Shortwave Radiation | Radiative exchange associated with shorter wavelengths that are emitted by the sun or reflected and absorbed by a surface<br>wavelength range includes UV to NIR |
| Normal-hemispherical | The method used to measure the reflection of light when a beam of light is incident on a surface, then measured with a detector that captures light in a hemisphere. |
| Normal-normal | The method used to measure the reflection of light when a beam of light is incident on a surface, then measured with a detector that captures light in a small solid angle close to the incident beam |
| Absorption | The process of a material taking in electromagnetic radiation (light) and converting it to internal energy |
| Absorptance | The fraction of incident radiation that is absorbed by a surface or material; ranges from 0 to 1. |
| Emittance | The fraction of blackbody radiation that is emitted by a surface or material for a given temperature; ranges from 0 to 1. |
| Reflectance | The fraction of incident radiation that is reflected by a surface or material; ranges from 0 to 1. |

**Table 2.**
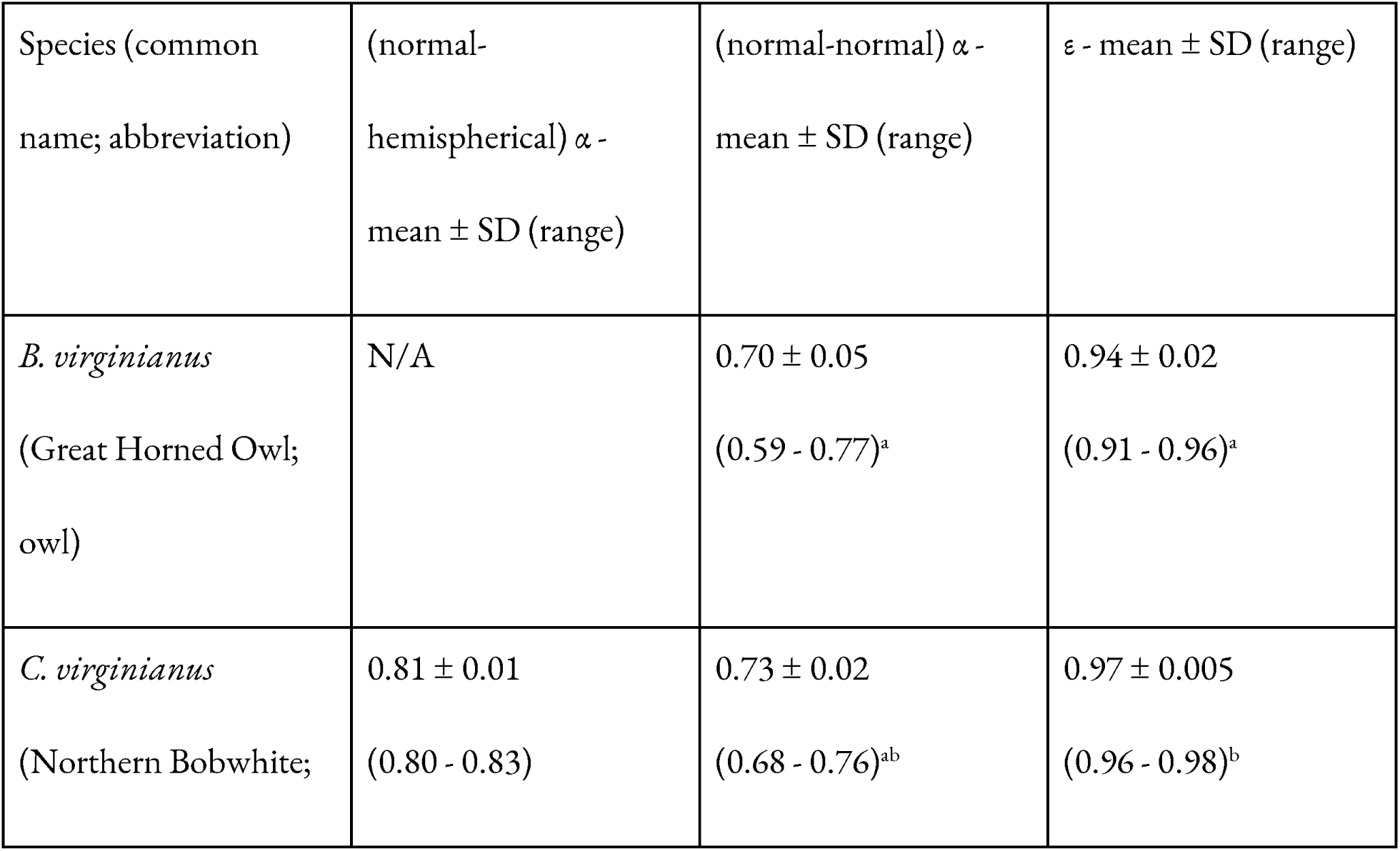
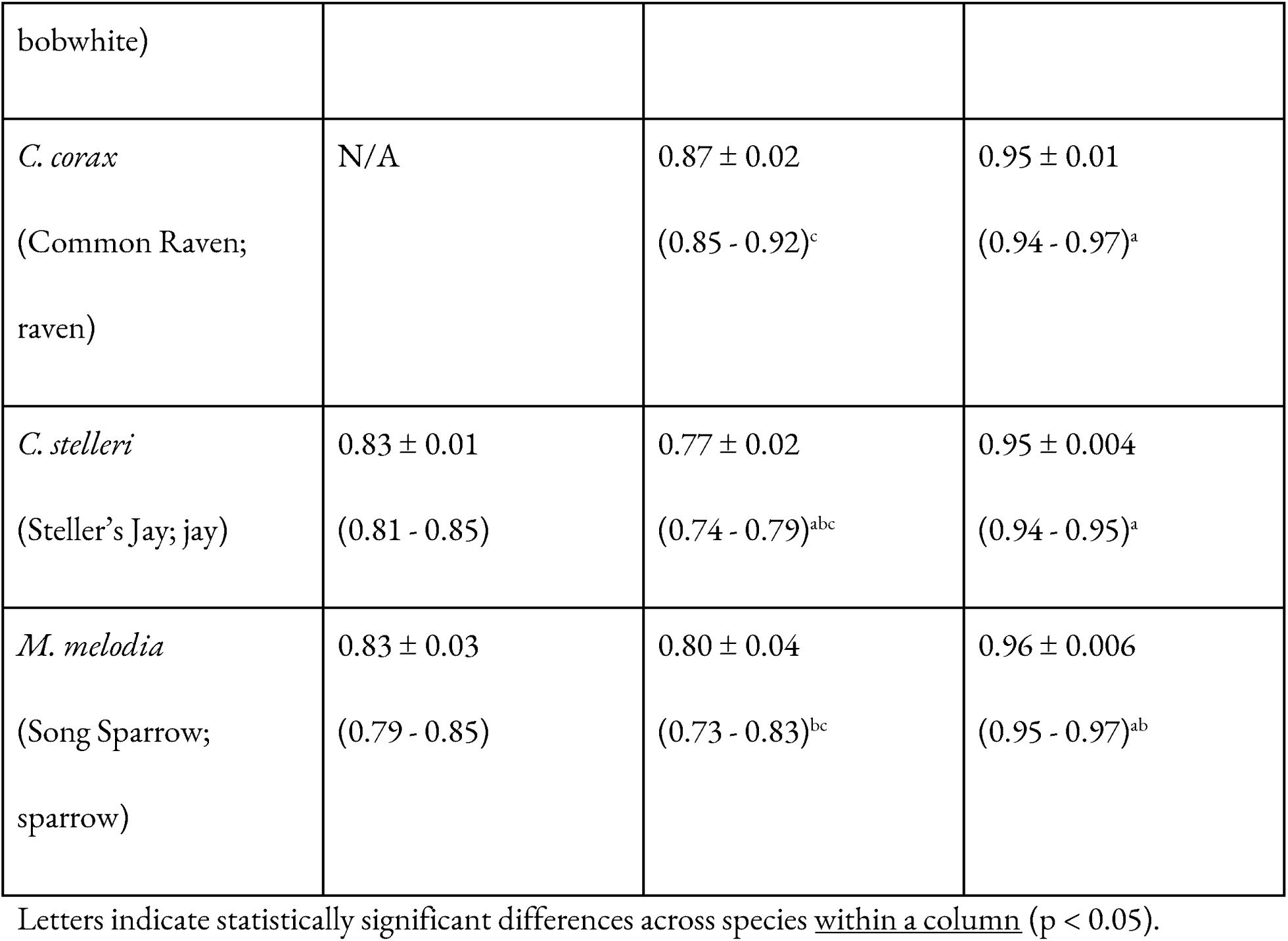
Mean absorption and emittance coefficients for each species.

There was also variation in mean total hemispherical emittance (ε) across species (Table 2, Figure 1; Kruskal-Wallis; KW = 24.50, p < 0.0001). Bobwhites had significantly higher emittance than all species except sparrows (Dunn’s MCT; bobwhite-owl: Z = 3.89, p = 0.001; bobwhite-raven: Z = 3.46, p = 0.0053; bobwhite-jay: Z = 4.44, p < 0.0001). All other species were not significantly different (Figures 1 and 2; Z < 2.48, p > 0.05).

In both cases, owls had the greatest range of absorptance and emittance, and thus the most variation among individuals. The range of absorptance in UV-NIR across species was much greater than the range of emittance in MIR (UV-NIR range of means: 0.70-0.87; MIR: 0.94-0.97; Table 2).

### Population-level trends in human-visible (400-700 nm) absorptance

Gloger’s rule has been based on differences in reflectivity across the human-visible spectrum (400-700 nm), not on the total solar spectrum (300-2500 nm). To assess if there were population-level trends in any of our focal species consistent with Gloger’s rule, we tested for human-visible changes in coloration (and bird-visible results [300-700 nm] are in Supplemental Table 2). Results were also analyzed as ‘average brilliance’ (where the reflectance at each wavelength from 400 - 700 nm is averaged, without accounting for incoming solar energy as in the absorption coefficient analysis), a technique commonly used for birds. However, population-level analyses of brilliance did not differ from the absorptance coefficient analysis (Supplemental Table 3), so we present only the absorptance coefficient analysis below.

There were significant differences among populations/subspecies in visible absorptance for jays and sparrows (Table 3; normal-hemispherical); in both cases, absorptance negatively correlated with the population’s mean annual temperature (linear regression; jay, F = 6.10, df = 7, p = 0.0428, [VIS absorptance] = -0.003[temperature] + 0.96; sparrow, F = 9.27, df = 7, p = 0.0382, [VIS absorptance] = -0.005[temperature] + 1.02). Jays from Mexico and California did not differ in their absorptance (Tukey’s MCT: q = 1.79, df = 6, p = 0.46), while jays from Alaska had increased absorptance (visibly darker coloration) compared to both other populations (Alaska-Mexico: q = 9.82, df = 6, p = 0.0011; Alaska-California: q = 8.03, df = 6, p = 0.0031).

**Table 3.** Differences in human-visible (400 - 700 nm) absorptance across populations.

| Species | Subspecies* | Populations (average temp) | $\alpha$ - mean ± SD (n) | P-value |
| --- | --- | --- | --- | --- |
| Owl** | *occidentalis* | Wyoming, USA/Canada (5 C) | 0.86 ± 0.04 (3) | F = 1.86; df =<br>2; p = 0.24 |
|  | *pallascens* | California, USA (desert) (15.5 C) | 0.82 ± 0.05 (3) |  |
|  | *pacificus* | California, USA (coast) (18.4 C) | 0.89 ± 0.009 (3) |  |
| Bobwhite | *mexicanus* | Iowa, USA (10.6 C) | 0.97 ± 0.009 (3) | F = 0.01; df =<br>2; p = 0.99 |
|  | *insignis* | Chiapas, MX (19.3 C) | 0.97 ± 0.003 (3) |  |
|  | *floridanus* | Florida, USA (24.3 C) | 0.97 ± 0.006 (3) |  |
| Raven** | *principalis* | Alaska, USA (8.4 C) | 0.99 ± 0.007 (3) | F = 0.47; df =<br>2; p = 0.65 |
|  | *sinatus* (A) | California, USA (16.4 C) | 0.99 ± 0.01 (3) |  |
|  | *sinatus* (B) | Sinaloa, MX/BajaCa, MX (24.2 C) | 0.99 ± 0.004 (3) |  |
| Jay | *stellari<sup>a</sup>* | Alaska, USA (8.2 C) | 0.99 ± 0.004 (3) | F = 27.35; df =<br>2; **p = 0.001** |
|  | *frontalis<sup>b</sup>* | California, USA (11.8 C) | 0.97 ± 0.006 (3) |  |
|  | *diademata<sup>b</sup>* | Chihuahua, MX (16.4 C) | 0.96 ± 0.004 (3) |  |
| Sparrow | *caurina<sup>a</sup>* | Alaska, USA (8.3 C) | 0.99 ± 0.005 (3) | K-W = 5.96; **p**<br>= **0.025** |
|  | *cooperi<sup>a</sup>* | California, USA (coastal) (16.7 C) | 0.98 ± 0.006 (3) |  |
|  | *saltonis<sup>a</sup>* | California, USA (desert) (18.4 C) | 0.96 ± 0.005 (3) |  |

\*letters indicate statistically significant differences using Tukey's posthoc test (following ANOVA) or
Dunn's posthoc test (following Kruskal-Wallis).
\*\*absorptance coefficients for owl and raven are normal-normal; the other species are normal-hemispherical.

Sparrows populations differed (K-W: 5.96, p = 0.025; normal-hemispherical). Sparrows from the deserts and coasts of California, and from Alaska and coastal California, did not differ in their absorptance coefficients (Alaska-coastal California: Z = 0.75, p > 0.99; coastal-desert California: Z = 1.64, p = 0.30). However, absorptance increased (darker coloration) in sparrows from Alaska compared to those from the desert region of California (Z = 2.39, p = 0.0512).

### Population-level trends in total solar absorptance (UV-NIR)

There were significant differences among populations/subspecies in solar absorptance for bobwhites and sparrows (ANOVA; Table 4; normal-hemispherical); populations with warmer mean temperatures had reduced absorptance compared to those with cooler temperatures. Bobwhites from Mexico and Florida did not differ in their absorptance coefficients (Tukey’s MCT: q = 2.32, df = 6, p = 0.30), while bobwhites from Iowa had increased absorptance (darker coloration) compared to both other populations (Figures 3 and 4; Iowa-Mexico: q = 8.01, df = 6, p = 0.0031; Florida-Iowa: q = 10.33, df = 6, p = 0.0008). Sparrows from Alaska and the coastal region of California did not differ in their absorptance coefficients (q = 0.26, df = 6, p = 0.98), while sparrows from the desert region of California had lower absorptance coefficients (lighter coloration) than both other populations (Figures 3 and 4; Alaska-desert California: q = 19.89, df = 6, p < 0.0001; coastal-desert California: q = 20.15, df = 6, p < 0.0001). In the case of bobwhites, absorptance was significantly and negatively correlated with mean annual temperature (F = 54.84, df = 7, p = 0.0001, [UV-VIS absorptance] = - 0.002[temperature] + 0.85), while there was a non-significant but negative trend in sparrows (F = 5.94, df = 4, p = 0.07).

**Figure 3.**
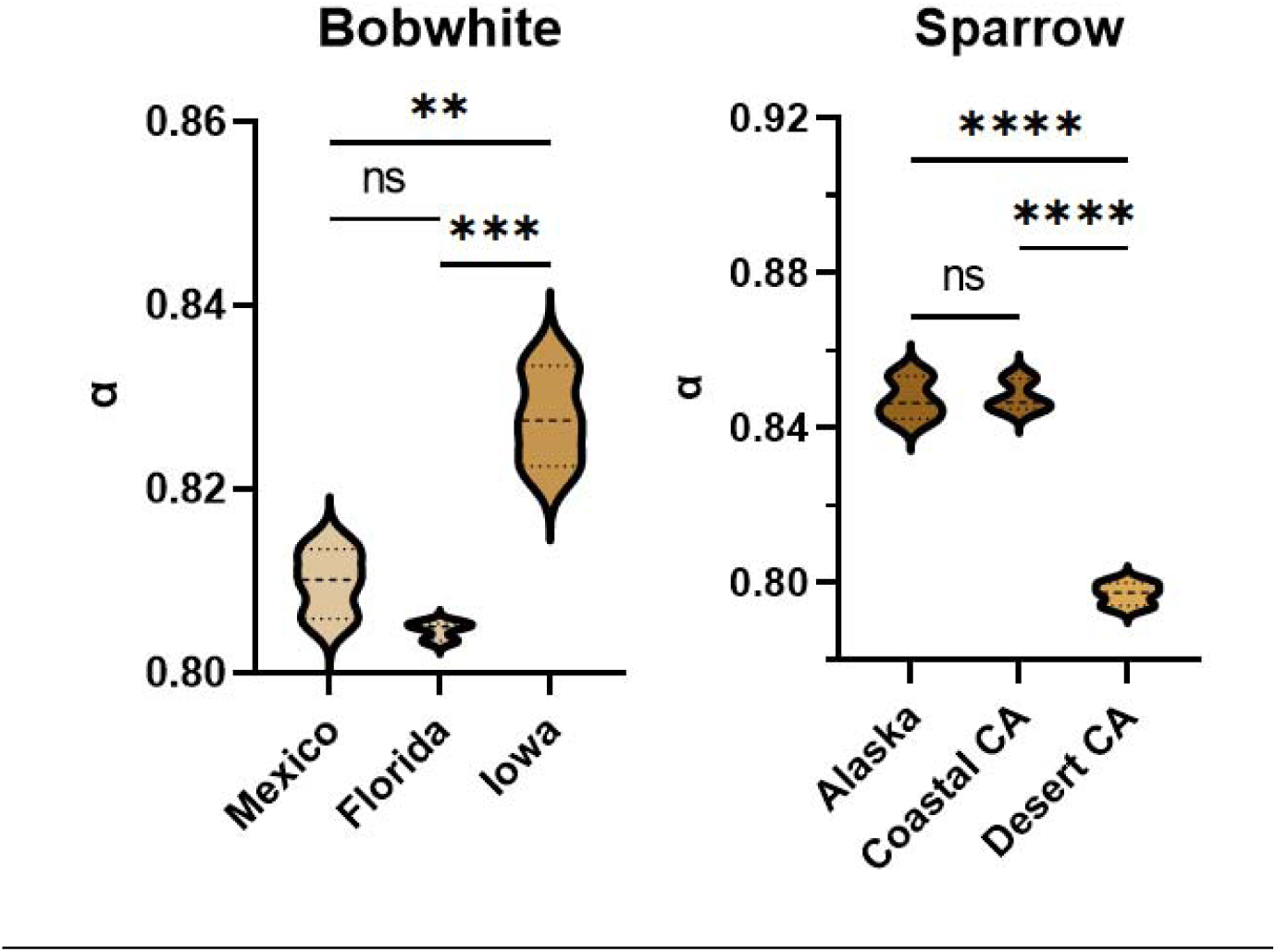
Population-level differences in solar absorptance coefficients for (left) bobwhites and (right) sparrows. Bobwhites from Iowa had increased absorptance compared to those from Mexico and Florida. Sparrows from the desert region of California had lower absorptance coefficients than those from Alaska and coastal California. ns = not significant; ** = p < 0.01; *** = p < 0.001; **** = p < 0.0001.

**Figure 4.**
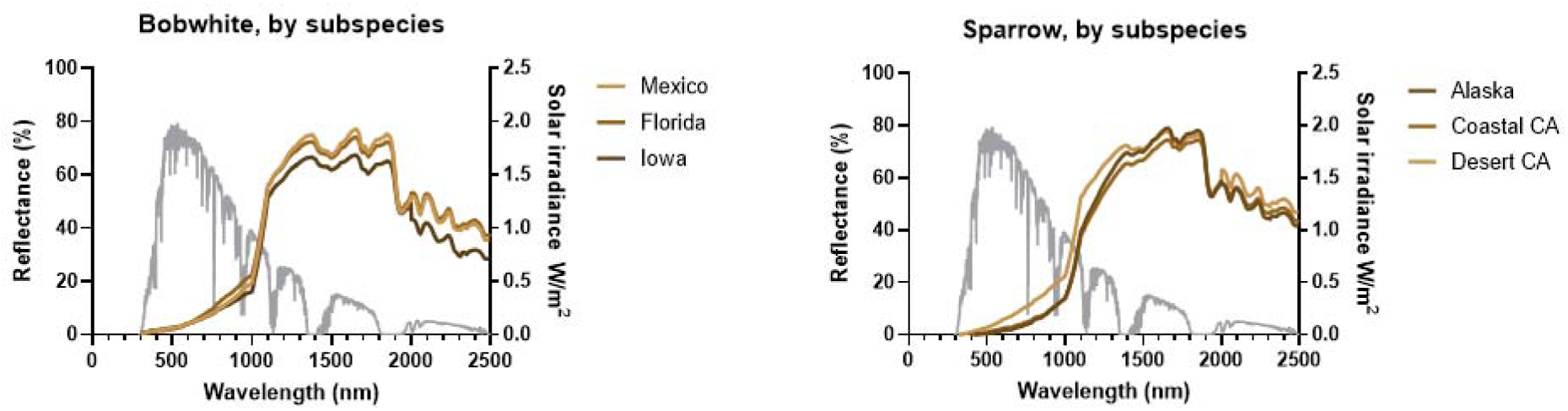
Population-level differences in reflectance across the solar spectra (300 - 2500 nm). Reflectance over the solar spectra is shown for each population with a solid line. The gray line shows solar irradiance in W/m^2^.

**Table 4.**
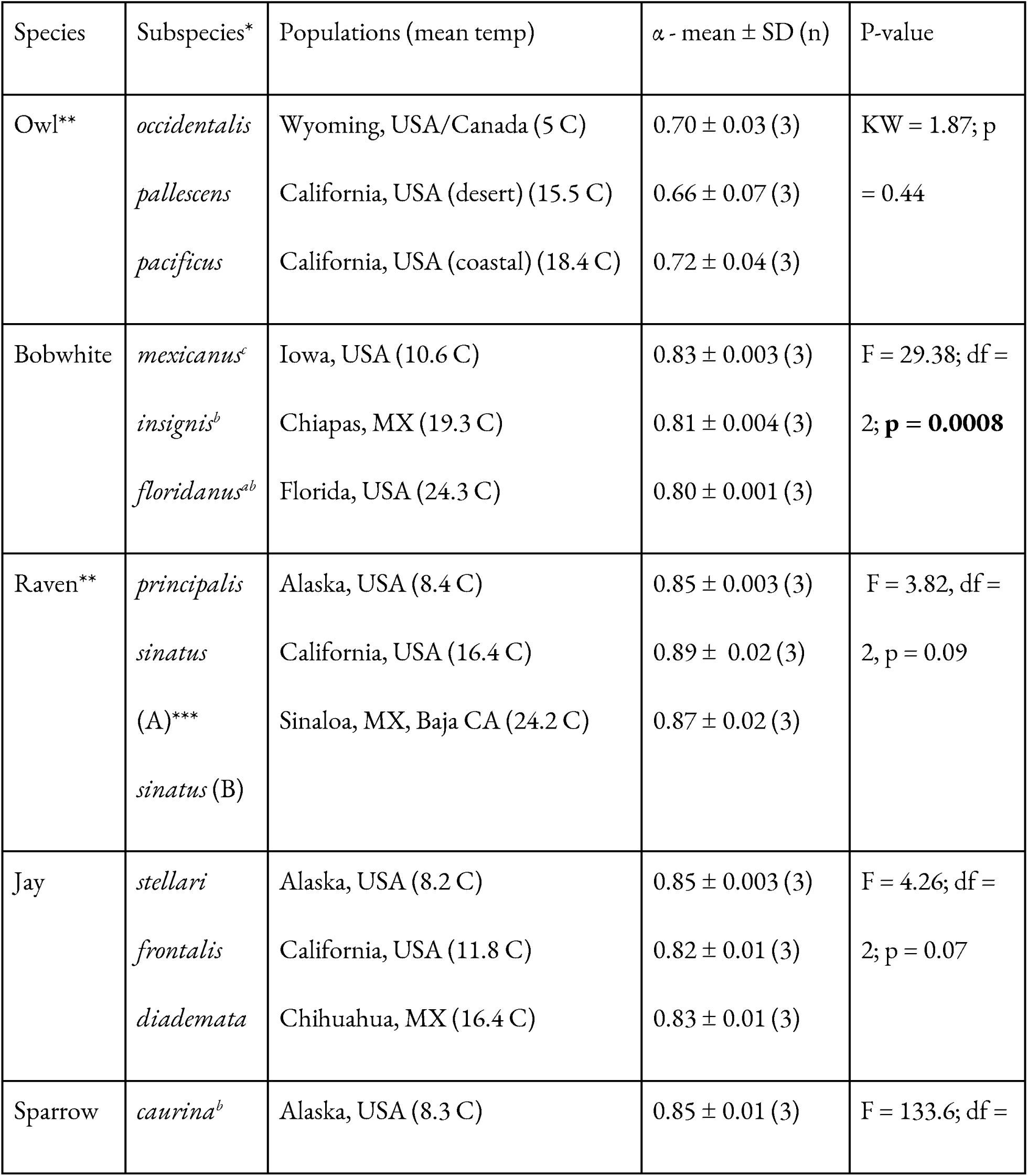
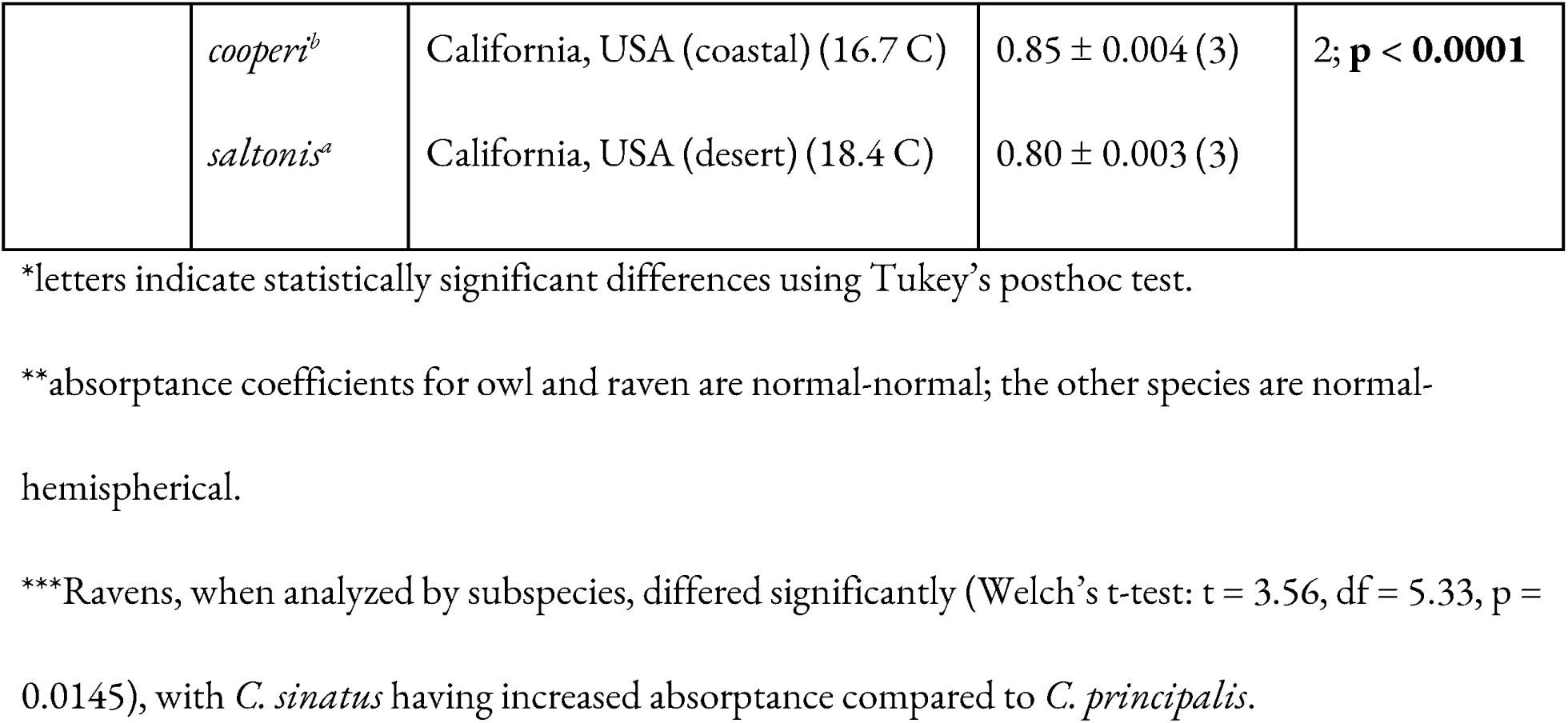
Differences in solar (UV-NIR) absorptance across populations.

Ravens did not show differences among populations (ANOVA; Table 4), but subspecies showed differences in absorptance (Welch’s t-test: t = 3.56, df = 5.33, p = 0.0145), where the mean absorptance of *C. corax sinatus* was significantly higher (darker coloration) than *C. corax principalis* (Supplemental Figure 2). In this case, the subspecies from the warmer locations had *increased* mean absorptance compared to the subspecies from the cooler location (the opposite pattern as seen in sparrows and bobwhites).

### Population-level trends in MIR emittance

There were significant differences among populations/subspecies in emittance for bobwhites (ANOVA; Table 5) but not owls, sparrows, ravens, or jays. Bobwhites from Iowa and Florida, and Florida and Mexico, did not differ in MIR emittance (Figure 5; Dunnett’s T3; Mexico-Florida: t = 0.03, df = 2.04, p > 0.99; Florida-Iowa, t = 2.32, df = 2.06, p = 0.29), while bobwhites from Iowa (the coolest population) had significantly higher emittance than those from Mexico (t = 15.25, df = 3.86, p = 0.0003). However, differences were not pronounced in the atmospheric transmission window (∼7.5 - 14 μm).

**Figure 5.**
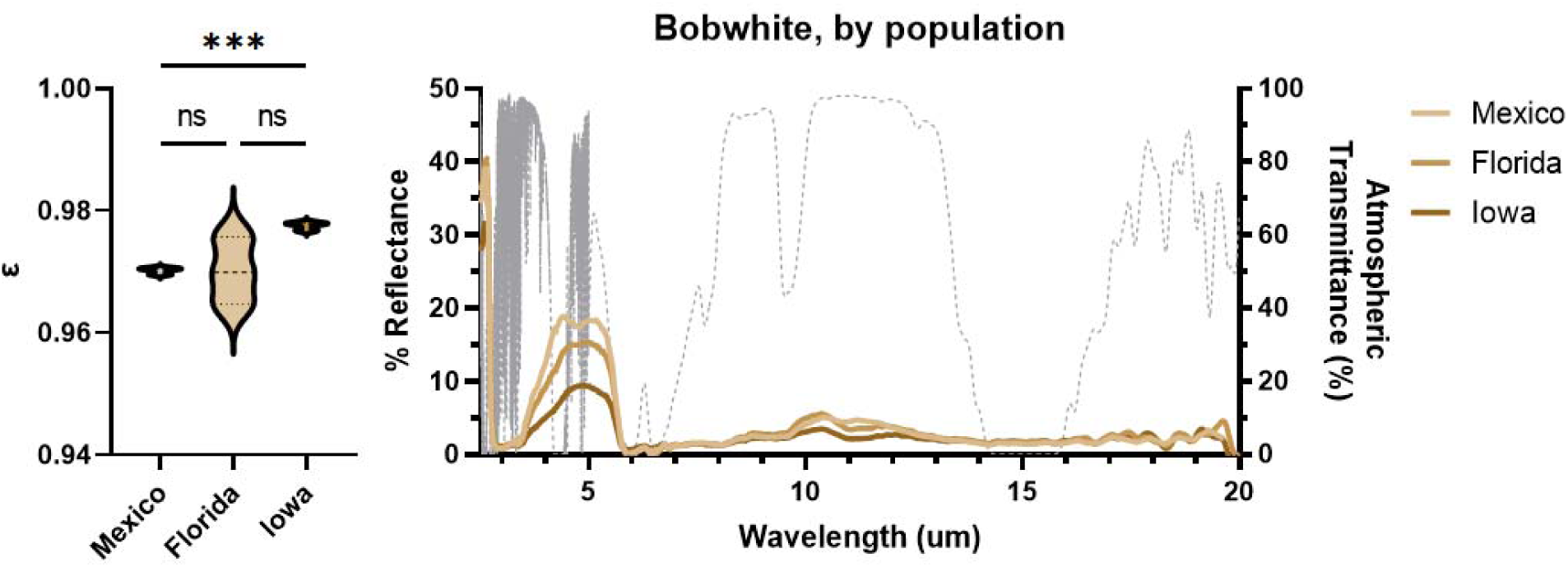
Population-level differences in (left) emittance coefficients and (right) MIR reflectance spectra in bobwhites. Bobwhites from Iowa and Florida, and Florida and Mexico, did not differ in their emittance coefficients, while bobwhites from Iowa had significantly higher emittance than those from Mexico. The gray dashed line shows atmospheric transmission (%). ns = not significant; *** = p < 0.001.

**Table 5.** Differences in emittance across populations of each species.

| Species | Subspecies | Populations (mean temp) | $\epsilon$ - mean ± SD (n) | P-value |
| --- | --- | --- | --- | --- |
| Owl | *occidentalis* | Wyoming, USA/Canada (5 C) | 0.94 ± 0.03 (3) | F = 0.24; df =<br>2; p = 0.79 |
| | *pallascens* | California, USA (desert) (15.5 C) | 0.94 ± 0.02 (3) | |
| | *pacificus* | California, USA (coastal) (18.4 C) | 0.95 ± 0.01 (3) | |
| Bobwhite | *mexicanus*<sup>b</sup> | Iowa, USA (10.6 C) | 0.98 ± 0.01 (3) | W = 98.23; df<br>= 2; p =<br>**0.0008\*** |
| | *insignis*<sup>a</sup> | Chiapas, MX (19.3 C) | 0.97 ± 0.001 (3) | |
| | *floridanus*<sup>ab</sup> | Florida, USA (24.3 C) | 0.97 ± 0.01 (3) | |
| Raven** | *principalis* | Alaska, USA (8.4 C) | 0.96 ± 0.01 (3) | F = 1.14, df =<br>2, p = 0.28 |
| | *sinatus* (A) | California, USA (16.4 C) | 0.95 ± 0.008 (3) | |
| | *sinatus* (B) | Sinaloa, MX, Baja CA (24.2 C) | 0.95 ± 0.006 (3) | |
| Jay | *stellari* | Alaska, USA (8.2 C) | 0.95 ± 0.004 (3) | F = 2.73; df =<br>2; p = 0.14 |
| | *frontalis* | California, USA (11.8 C) | 0.95 ± 0.003 (3) | |
| | *diademata* | Chihuahua, MX (16.4 C) | 0.95 ± 0.001 (3) | |
| Sparrow | *caurina* | Alaska, USA (8.3 C) | 0.96 ± 0.004 (3) | F = 1.37; df =<br>2; p = 0.32 |
| | *cooperi* | California, USA (coastal) (16.7 C) | 0.96 ± 0.005 (3) | |
| | *saltonis* | California, USA (desert) (18.4 C) | 0.96 ± 0.004 (3) | |

\*letters indicate statistically significant differences using Dunnett's T3 MCT.
\*\* no difference between subspecies: Unpaired t-test, $t = 1.15$ , $df = 7$ , $p = 0.29$

## Discussion

We found variation in UV-NIR absorptance and MIR emittance across species, and within some species across populations. For instance, owls have significantly lower mean absorptance in the UV-NIR than several diurnal species, such as ravens and sparrows, as well as significantly greater interindividual variation in reflectance across the UV-MIR compared to all other species. This increased variance may reflect relaxed selective pressure on reflectance of solar radiation across the entire solar spectrum in this species, as owls are nocturnal. Alternatively, owls are large and patterned (at least, in the VIS spectrum), with many layers of feathers; these factors may contribute to greater variance in measurement consistency among individuals. Reduced selective pressure on reflectance as a thermal adaptation may also explain why owl populations did not differ from one another in either UV-NIR absorptance or MIR emittance, despite being known to follow Gloger’s rule. We also found that bobwhites had notably higher MIR emittance than all other tested species, though interspecific differences in emittance were all within 3% for our tested species.

Our data indicate intraspecific differences across populations in mean absorptance in the UV-NIR for bobwhites and sparrows, and across subspecies in ravens. We observed a non-significant trend towards reduced overall UV-NIR absorptance in jays from warmer populations (consistent with a thermal hypothesis). Interestingly, bobwhites and sparrows both followed an inverse of Gloger’s rule in the UV-NIR, with darker bobwhites (e.g., increased absorptance) found in cooler regions/higher latitudes, and the same for sparrows. This suggests that these population-level differences in UV-NIR reflectance may be consistent with a thermally-driven hypothesis, wherein hotter environments may be populated by lighter colored (e.g., more reflective) individuals to reduce heat stress. Similar results were found in interspecific bird pigmentation analyses (Galván et al. 2018) and UV-NIR analyses (Medina et al. 2018), and have also been shown in gastropods and butterflies (Munro et al. 2019; Franklin et al. 2022).

Prior research has demonstrated that there are trade-offs between camouflage and thermoregulation, where brighter horned larks in a given environment had increased cooling but worse camouflage in the VIS (Mason et al. 2023). While we found that sparrow populations differed whether considering just the VIS or the entire UV-NIR, populations of bobwhites only differed in absorptance across the entire UV-NIR spectrum and not when only the VIS was considered. It is possible that decoupling the degree of variation in the VIS from the NIR allows bobwhites to achieve ‘the best of both worlds’ - remaining well-camouflaged in the VIS while reducing their heat load through changes in NIR/total solar absorptance. Bobwhites are open habitat grassland species that experience high levels of solar radiation depending on their habitat structure, potentially driving these stronger differences across populations.

Ravens are also open habitat nesters, but did not demonstrate decreased UV-NIR absorption among their populations - though when analyzed by subspecies, ravens in warmer climates actually had *increased* absorptance compared to those from cooler areas (the opposite of the pattern observed for sparrows and bobwhites). Although initially it would seem as though darker plumage must always increase body temperature through increased absorption of solar energy, studies in darkly colored birds have shown that darker plumage can sometimes offer a thermoregulatory advantage in environments with high solar load and high wind velocities, such as those produced by active flight (Wolf and Walsberg 2000). Lighter plumage can reflect more radiation, but also allows increased penetration of that radiation into the plumage, potentially resulting in increased heating of the skin; darker plumage absorbs more solar radiation, but this heat is kept at the surface of the animal where it can more easily be dispersed convectively (Walsberg 1982). Studies in the brown-necked raven showed that black plumage heated up more than light plumage at the animal’s surface, but skin temperature remained the same (Marder 1973); at wind velocities greater than 5.5 m/s, pigeons with black plumage had reduced solar heat loads compared to those with white plumage (Walsberg et al. 1978; Wolf and Walsberg 2000). Accordingly, ravens in warmer climates with increased solar load may benefit from the increased UV-NIR absorptance of their dorsal feathers by using convection to shed heat from their surface.

MIR emittance only differed by population in bobwhites, with greater dorsal emittance found in higher latitude populations. This is the first demonstration of intraspecific, population-level differences in MIR emittance in animals. The population differences in bobwhites were surprising in light of prior findings on insects by Krishna et al. (2021), as selection on MIR emittance should be only based on thermal pressures, which we expect to select for the opposite result (greater emittance on the dorsal surface in hotter climates to offload heat through the atmospheric transmission window; see, ants: Shi et al. 2015). There are several possible explanations to explain why we found the opposite of what we might have expected: 1) Cloud coverage and atmospheric moisture might be better predictors of population-level variance in MIR emittance than temperature or latitude, as these variables impact the strength of the sky as a radiative sink (see Hardy and Stoll 1954 v. Nobel 1991) and thus could be more important than air temperature in determining how the animal’s energy balance is impacted by emittance (though see Krishna et al. 2021, where precipitation was not linked to MIR emissivity in butterflies); 2) that emittance varies based on the ‘side’ of the feather measured (e.g., emittance of MIR radiation from the bird’s body going into the environment may differ from emittance of external MIR radiation entering the bird’s body); and/or 3) that differences in MIR emittance are so slight among populations as to be negligible to the animals’ heat budget, resulting in largely no patterns among populations with occasional statistically significant ‘noise’. Given that differences in emissivity were only 1% across populations, and that longwave radiative heat exchange generally represents a relatively smaller fraction of birds’ energy budgets (e.g., ∼8%, Ward et al. 1999; though see Léger and Larochelle 2006 under more naturalistic conditions), it seems plausible that if hypothesis two can be ruled out, hypothesis three is most likely in the species of birds we tested.

While conducting this study, we encountered some methodological constraints that deserve further attention in studies that focus on UV-NIR absorptance and MIR emittance in birds. For example, some birds have clearer visible patterning on the dorsal surface than others (consider owls vs. ravens); accounting for this variance when using spectrometers that only assess a ‘spot’ of a certain diameter can be challenging in the VIS range. This is less likely to impact NIR and MIR analyses, where this patterning generally becomes less relevant or may even disappear (though localized MIR emittance differences are known in ants and butterflies; Shi et al. 2015; Krishna et al. 2020); and localized UV-NIR differences have also been shown in lizards Smith et al. 2016; Barrett and O’Donnell 2023). Further, the size of birds can create challenges when attempting to obtain measurements - most spectrometers are designed for use with small material samples, not large specimens like many birds. In our study, this constrained our ability to obtain normal-hemispherical NIR absorptance for all species (which would be considered the more ecologically relevant measure). Owls and ravens were too large to fit in the spectrometer, and only normal-normal absorptance could be obtained. Further, analyzing feathers separated from birds did not provide accurate results (presumably due to a lack of appropriate layering), with a 3-fold difference in measured dorsal reflectance for an intact song sparrow compared to a single feather (Supplemental Figure 3).

For species where both normal-normal and normal-hemispherical data could be obtained, means differed by 3 - 8% between the measurements, suggesting that study results could differ based on the ability to obtain normal-hemispherical vs. normal-normal data on whole birds (Table 1).

Feather morphology may magnify the different relationships between normal-hemispherical and normal-normal data across species, with some feathers likely to increase the amount of diffuse reflectance off the sample - thus increasing the relevance of normal-hemispherical data.

Our data reinforce prior findings that NIR absorptance varies among birds, and extends our understanding to intraspecific variation corresponding to habitat characteristics. This variation correlates with latitude in a manner opposite with the predictions of Gloger’s rule - where birds in warmer climates, or at lower latitudes, tend to have reduced absorptance (e.g., lighter color). This suggests that warmer climates and increased solar load may drive the evolution of reduced NIR absorptance in birds’ dorsal feathers as a thermal adaptation. Further, our results suggest detectable differences in MIR emittance across populations in some species. However, these differences in MIR emittance are quite slight in the species we tested (no greater than 5% in either interspecific or intraspecific variation), representing very little of the animal’s overall thermal budget, and thus suggesting that thermal selective pressure on MIR emittance is not as great in endotherms as has been found in some ectotherms (variation in emissivity of 40% in butterflies from different climates; Krishna et al. 2021). In this study, we intentionally chose non-migratory birds, however an expanded study that considered additional life history characteristics (such as migration) may find greater differences, as some nocturnal migratory birds could be expected to have a greater reliance on longwave radiative heat exchange through the atmospheric transmission window (Léger and Larochelle 2006).

## Supplemental Materials

**Supplemental Figure 1.**
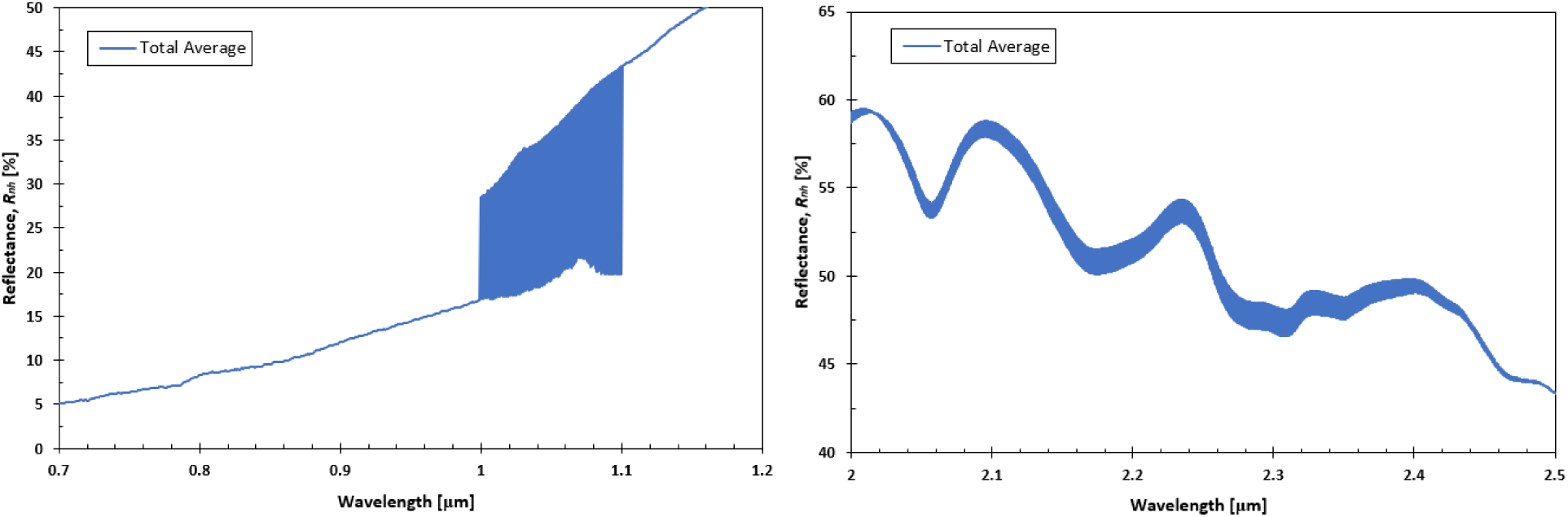
Difference in total average reflectance for sparrows (n = 9) based on detector type. **(left)** For the region between 1 and 1.1 μm, data sets were collected using both UV-Vis and InGaAs detectors; there was a difference in reflectance dependent on the type of detector used (difference demonstrated by the shaded region). As reflectance was approximately linear in this region, a linear interpolation was used to merge both data sets. **(right)** For the region between 2 and 2.5 μm, the MCT and InGaAs detectors had similar curves with slightly dissimilar values for total reflectance (difference demonstrated by the shaded region). MCT data was used so that the entire spectrum from 2 to 20 μm was collected in the same fashion, with no need for data manipulation.

**Supplemental Figure 2.**
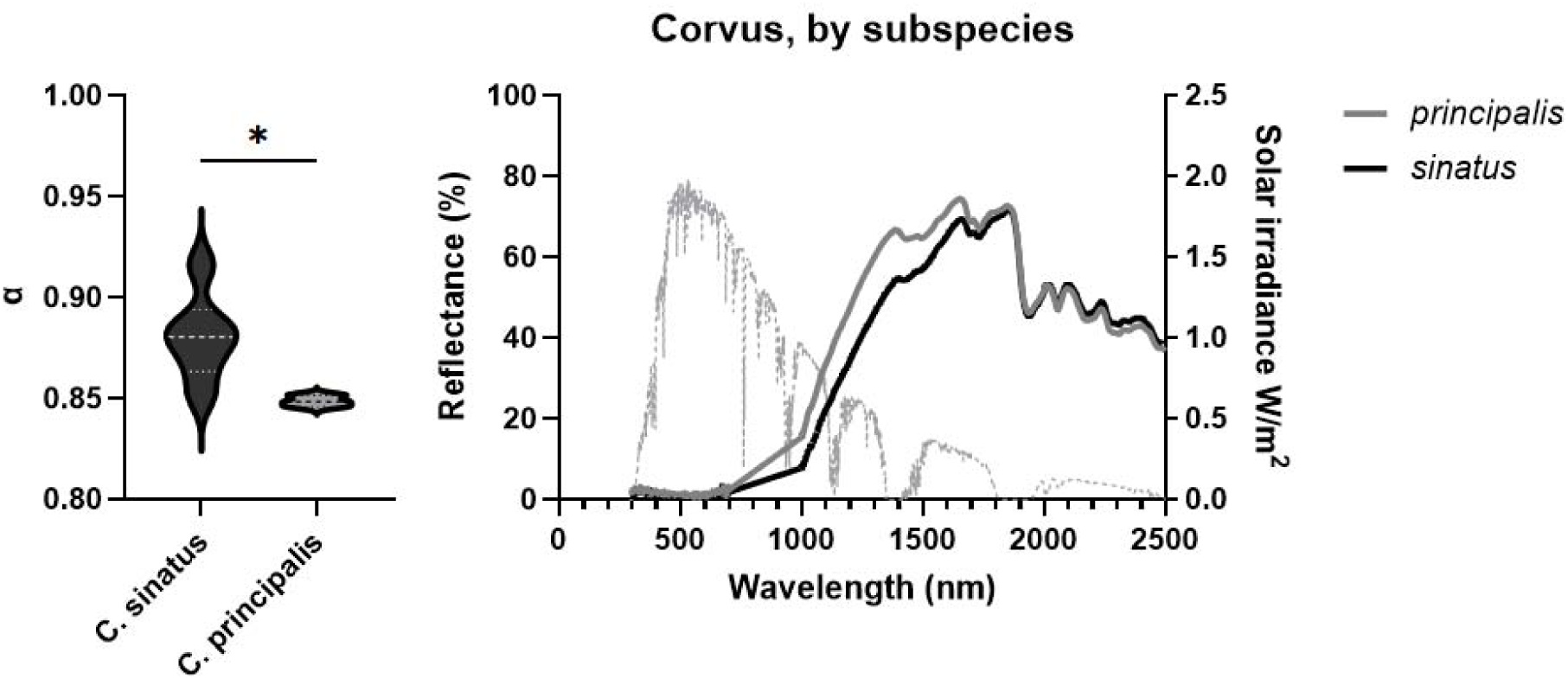
Subspecies-level (left) differences in total absorption coefficients and (right) reflectance spectra for ravens. **(left)** *C. corax sinatus* had significantly higher absorption than *C. corax principalis* (Welch’s t-test: t = 3.56, df = 5.33, p = 0.0145). **(right)**. Reflectance over the solar spectra is shown for each subspecies with a solid line. The gray dashed line shows solar irradiance in W/m^2^. * = p < 0.05

**Supplemental Figure 3.**
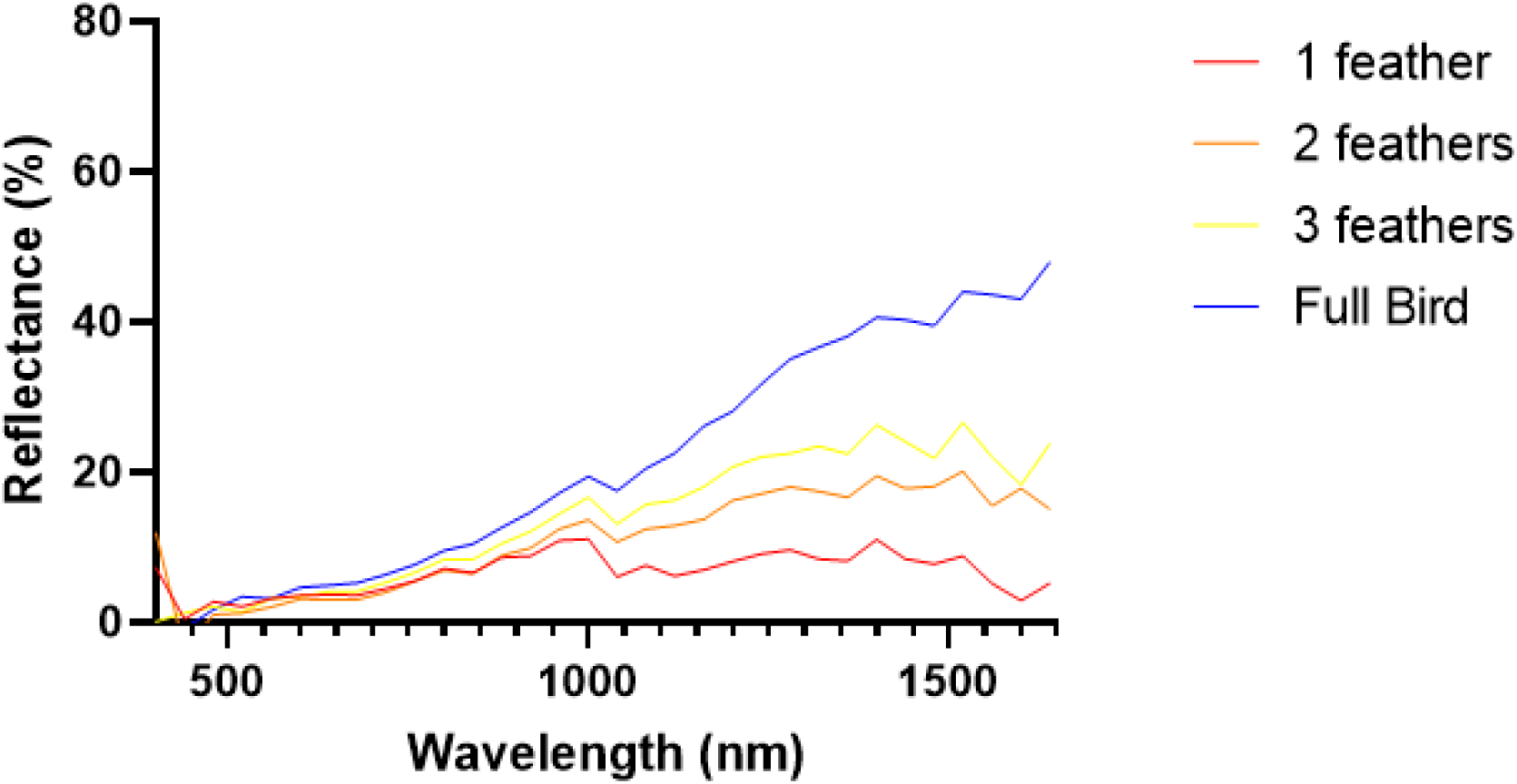
Difference in VIS-NIR reflectance for stacked dorsal feathers separated from a full bird v. the full bird (song sparrow). Average reflectance from 400 - 1600 nm was 6.87% for a single feather, 10.69% for two feathers, 13.75% for three feathers, and 20.35% for a dorsal surface measurement of the full bird, demonstrating the critical importance of correct feather stacking.

**Supplemental Table 1.**
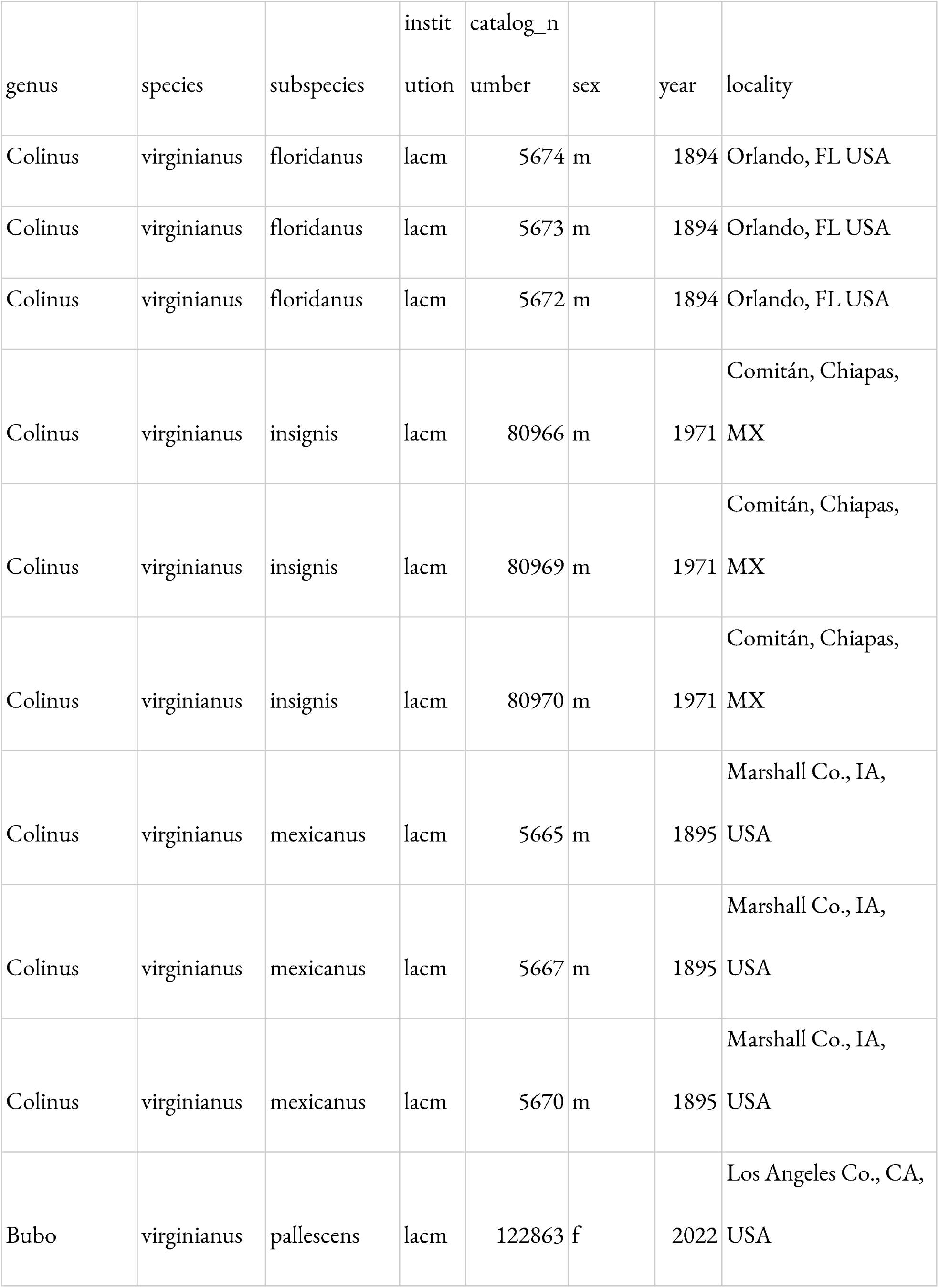
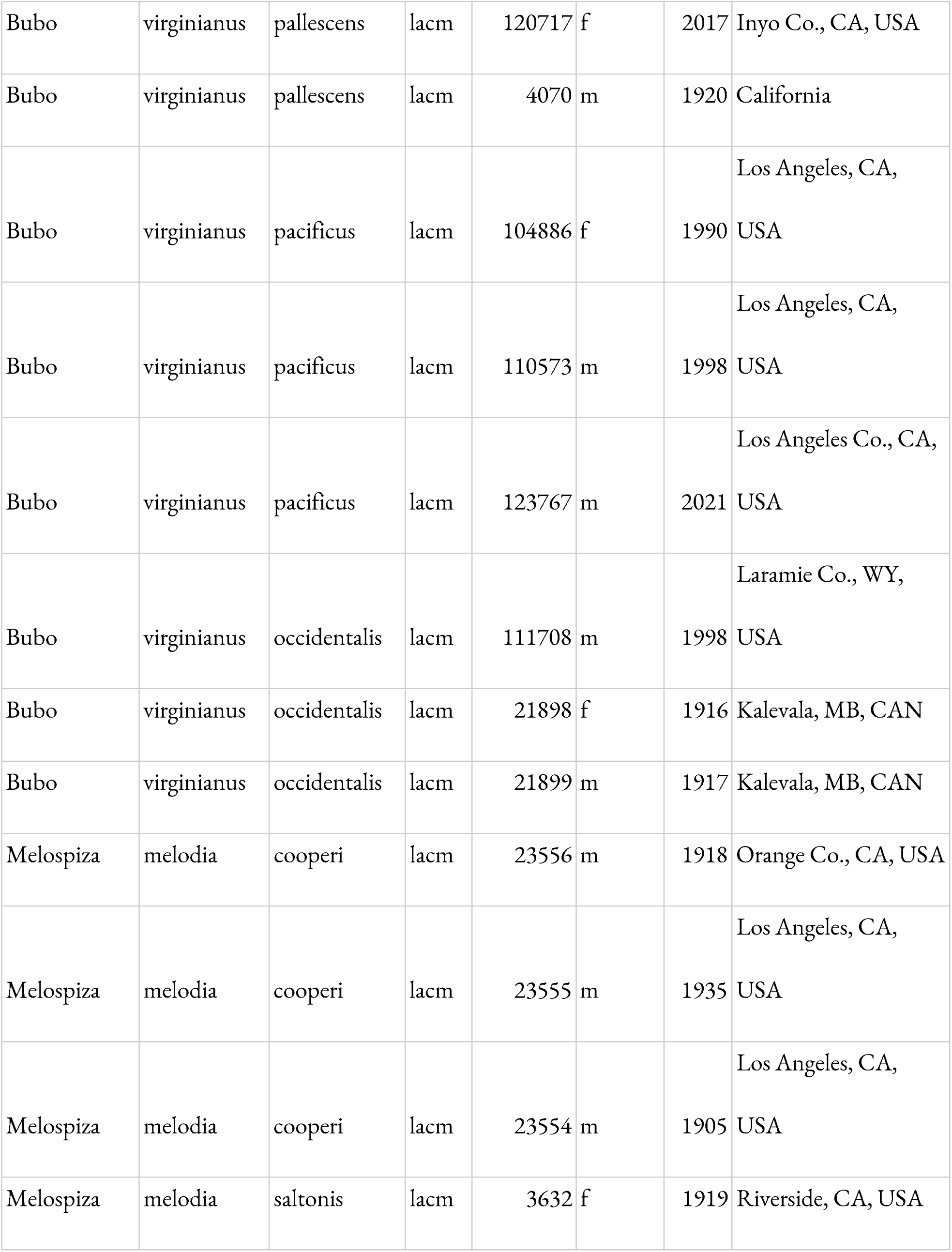
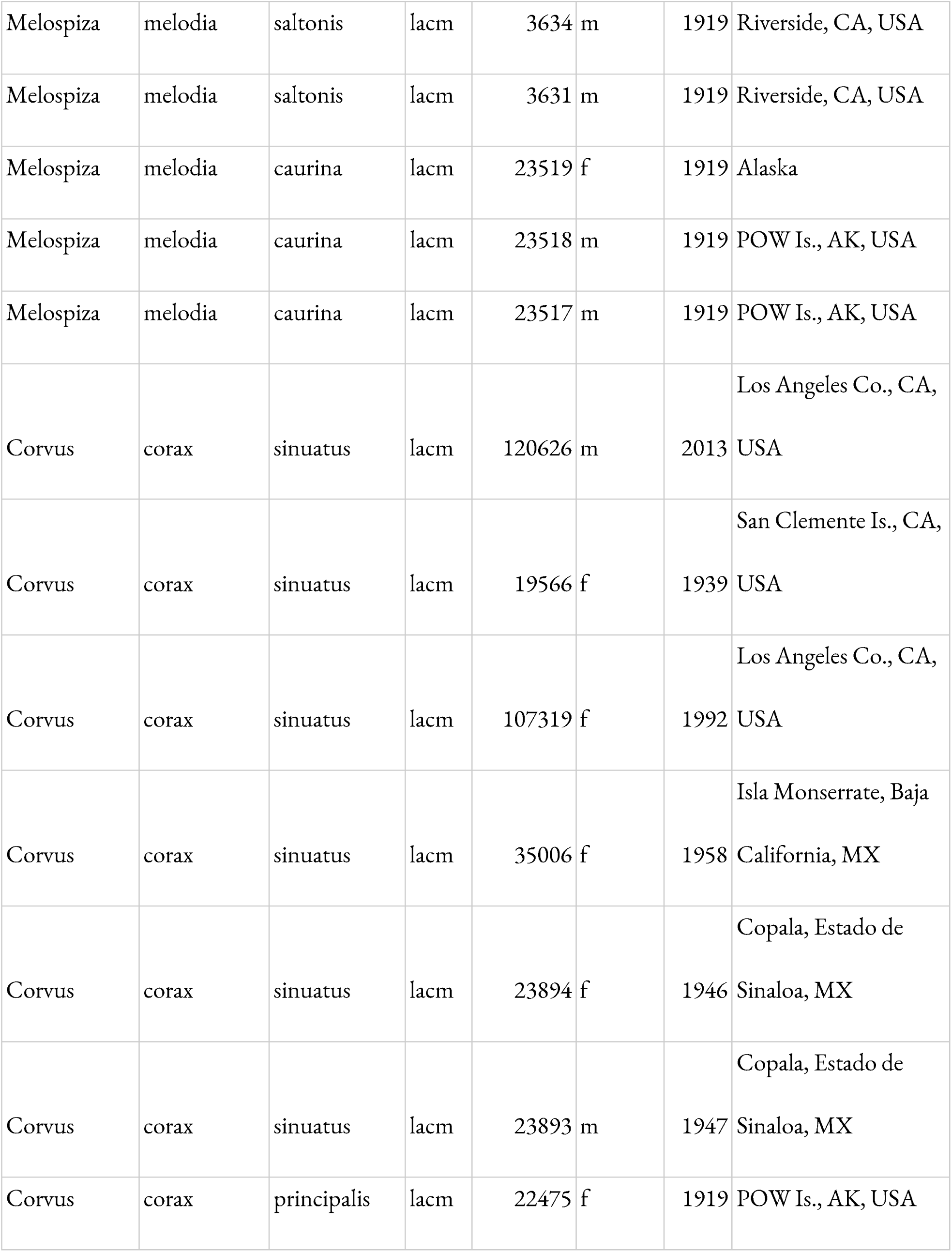
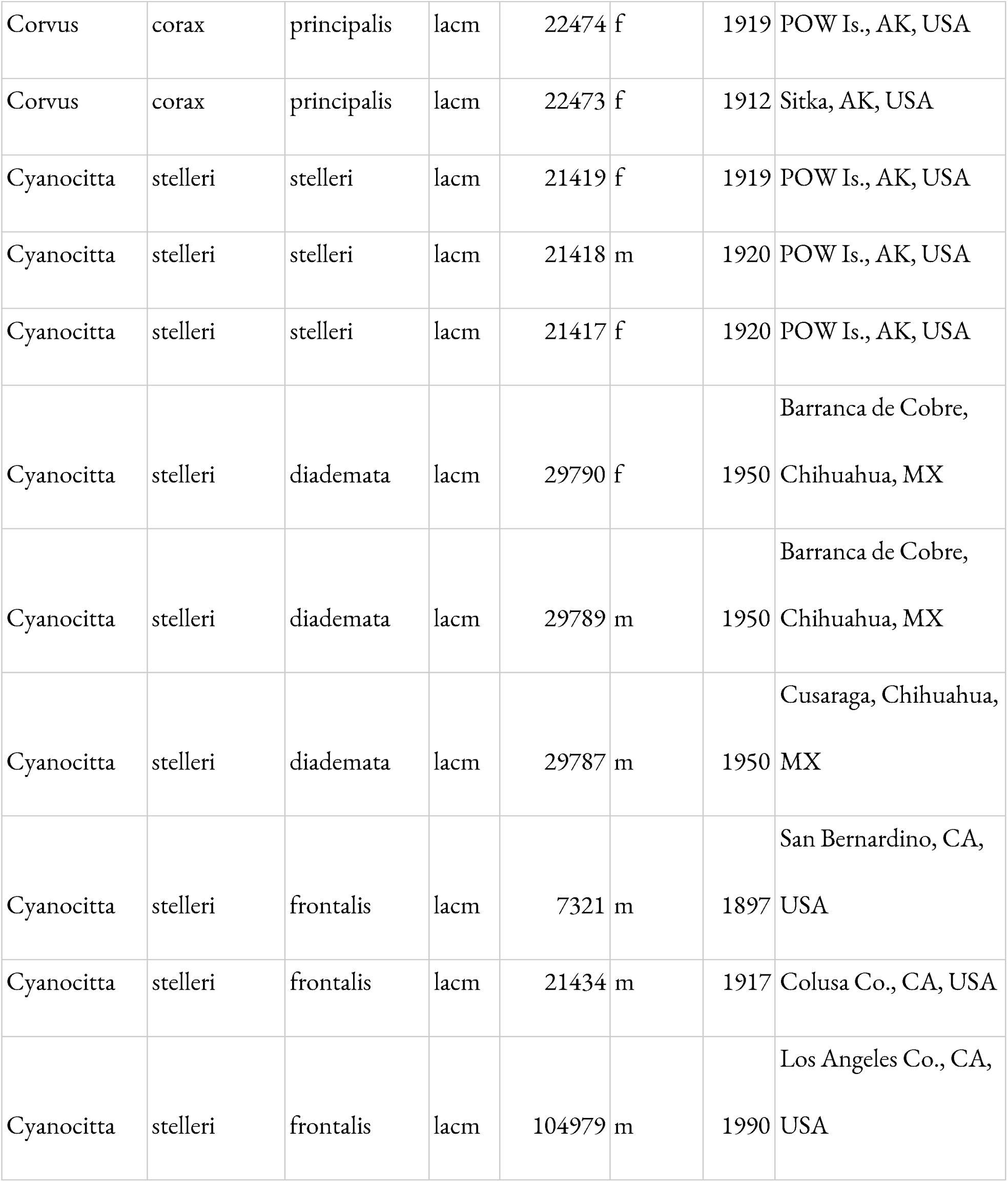
Catalog numbers and details of each specimen.

**Supplemental Table 2.**
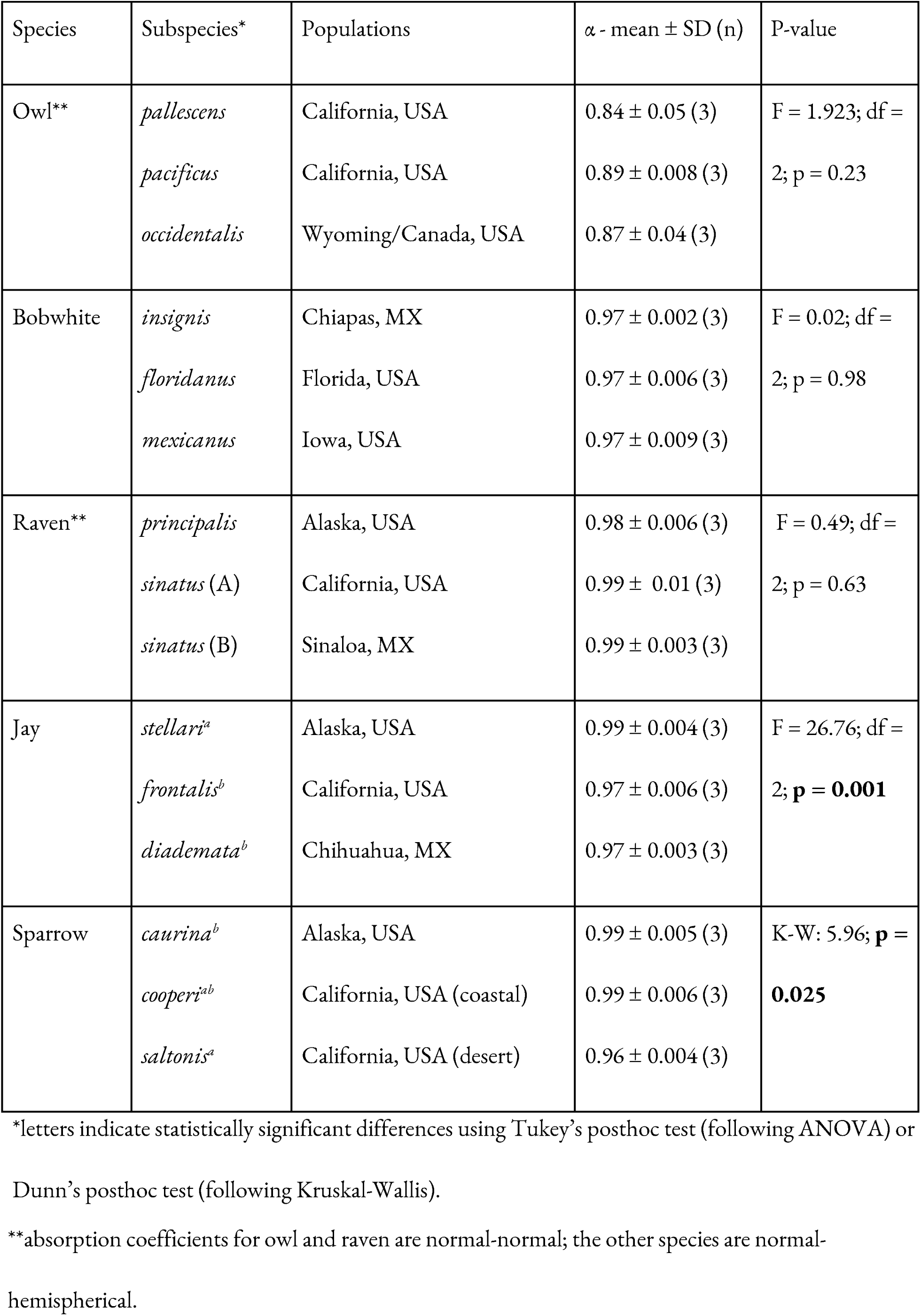
Differences in bird-visible (300 - 700 nm) absorption across populations.

| Species | Subspecies* | Populations | $\alpha$ - mean ± SD (n) | P-value |
| --- | --- | --- | --- | --- |
| Owl** | *pallescentis* | California, USA | 0.84 ± 0.05 (3) | F = 1.923; df =<br>2; p = 0.23 |
| | *pacificus* | California, USA | 0.89 ± 0.008 (3) | |
| | *occidentalis* | Wyoming/Canada, USA | 0.87 ± 0.04 (3) | |
| Bobwhite | *insignis* | Chiapas, MX | 0.97 ± 0.002 (3) | F = 0.02; df =<br>2; p = 0.98 |
| | *floridanus* | Florida, USA | 0.97 ± 0.006 (3) | |
| | *mexicanus* | Iowa, USA | 0.97 ± 0.009 (3) | |
| Raven** | *principalis* | Alaska, USA | 0.98 ± 0.006 (3) | F = 0.49; df =<br>2; p = 0.63 |
| | *sinatus* (A) | California, USA | 0.99 ± 0.01 (3) | |
| | *sinatus* (B) | Sinaloa, MX | 0.99 ± 0.003 (3) | |
| Jay | *stellari*<sup>a</sup> | Alaska, USA | 0.99 ± 0.004 (3) | F = 26.76; df =<br>2; **p = 0.001** |
| | *frontalis*<sup>b</sup> | California, USA | 0.97 ± 0.006 (3) | |
| | *diademata*<sup>b</sup> | Chihuahua, MX | 0.97 ± 0.003 (3) | |
| Sparrow | *caurina*<sup>b</sup> | Alaska, USA | 0.99 ± 0.005 (3) | K-W: 5.96; **p = 0.025** |
| | *cooperi*<sup>ab</sup> | California, USA (coastal) | 0.99 ± 0.006 (3) | |
| | *saltonis*<sup>a</sup> | California, USA (desert) | 0.96 ± 0.004 (3) | |

\*letters indicate statistically significant differences using Tukey's posthoc test (following ANOVA) or Dunn's posthoc test (following Kruskal-Wallis).
\*\*absorption coefficients for owl and raven are normal-normal; the other species are normal- hemispherical.

**Table 3.** Differences in average brilliance (400 - 700 nm) across populations.

| Species | Subspecies* | Populations | Brilliance (%) -<br>mean ± SD (n) | P-value |
| --- | --- | --- | --- | --- |
| Owl** | *pallescentis* | California, USA | 14.98 ± 4.44 (3) | F = 1.91; df =<br>2; p = 0.23 |
| | *pacificus* | California, USA | 9.81 ± 0.66 (3) | |
| | *occidentalis* | Wyoming/Canada, USA | 11.71 ± 3.47 (3) | |
| Bobwhite | *insignis* | Chiapas, MX | 2.86 ± 0.18 (3) | F = 0.03; df =<br>2; p = 0.97 |
| | *floridanus* | Florida, USA | 2.81 ± 0.51 (3) | |
| | *mexicanus* | Iowa, USA | 2.93 ± 0.80 (3) | |
| Raven** | *principalis* | Alaska, USA | 1.64 ± 0.64 (3) | F = 0.55; df =<br>2; p = 0.61 |
| | *sinatus* (A) | California, USA | 1.08 ± 1.00 (3) | |
| | *sinatus* (B) | Sinaloa, MX | 1.54 ± 0.31 (3) | |
| Jay | *stellari*<sup>a</sup> | Alaska, USA | 0.87 ± 0.40 (3) | F = 25.78; df =<br>2; **p = 0.0011** |
| | *frontalis*<sup>b</sup> | California, USA | 2.70 ± 0.50 (3) | |
| | *diademata*<sup>b</sup> | Chihuahua, MX | 3.13 ± 0.30 (3) | |
| Sparrow | *caurina*<sup>a</sup> | Alaska, USA | 0.67 ± 0.45 (3) | K-W = 5.96; **p** |
|  | *cooperi*<sup>a</sup> | California, USA (coastal) | 1.15 ± 0.56 (3) | = **0.025** |
|  | *saltonis*<sup>a</sup> | California, USA (desert) | 3.31 ± 0.36 (3) |  |

\*letters indicate statistically significant differences using Tukey's posthoc test (following ANOVA) or
Dunn's posthoc test (following Kruskal-Wallis).
\*\*absorption coefficients for owl and raven are normal-normal; the other species are normal-
hemispherical.

